# Structural and functional analysis of natural capsid variants reveals sialic-acid independent entry of BK polyomavirus

**DOI:** 10.1101/2022.07.13.499703

**Authors:** M.N. Sorin, A. Di Maio, L.M. Silva, D. Ebert, C. Delannoy, N.-K. Nguyen, Y. Guerardel, W. Chai, F. Halary, K. Renaudin-Autain, Y. Liu, C. Bressollette-Bodin, T. Stehle, D. McIlroy

**Affiliations:** Nantes Université, CHU Nantes, INSERM, Center for Research in Transplantation and Translational Immunology, UMR 1064, F-44000 Nantes, France; Interfaculty Institute of Biochemistry, University of Tübingen, Germany; Glycoscience Laboratory, Department of Metabolism, Digestion and Reproduction, Imperial College London, London, United Kingdom; Université de Lille, CNRS, UMR 8576 – UGSF - Unité de Glycobiologie Structurale et Fonctionnelle, F-59000 Lille, France; Institute for Glyco-core Research (iGCORE), Gifu University, Gifu, Japan; CHU Nantes Service d’Anatomie et Cytologie Pathologique, Medical, Nantes, France; CHU Nantes Laboratoire de Virologie, Nantes, France; Faculté de Médecine, Nantes Université, Nantes, France; Faculté des Sciences et des Techniques, Nantes Université, Nantes, France

## Abstract

BK Polyomavirus (BKPyV) is an opportunistic pathogen that causes nephropathy in kidney transplant recipients. The BKPyV major capsid protein, VP1, engages gangliosides, lipid-linked sialylated glycans at the cell surface, to gain entry into cells. Here, we characterise the influence of VP1 mutations observed in patients with persistent post-transplant BKPyV replication on ganglioside binding, VP1 protein structure, and the tropism of the virus in two renal cell lines: 293TT and immortalised renal tubular epithelial (RS) cells. Infectious entry of single mutants E73Q, E73A and the triple mutant A72V-E73Q-E82Q (VQQ) remained sialic acid-dependent. These three variants acquired binding to a-series gangliosides, including GD1a, although only E73Q was able to infect GD1a-supplemented LNCaP or GM95 cells. Crystal structures of the three mutants showed a clear shift of the BC2 loop in mutants E73A and VQQ that correlated with the inability of these VP1 variants to infect ganglioside complemented cells. On the other hand, the double mutant K69N-E82Q lost the ability to bind sialic acid, with the K69N mutation leading to a steric clash which precludes sialic acid binding. Nevertheless, this mutant retained significant infectivity in 293TT cells that was not dependent on heparan sulphate proteoglycans, implying that an unknown sialic acid-independent entry receptor for BKPyV exists.

## Introduction

BK Polyomavirus (BKPyV) is a small non-enveloped double stranded DNA virus with an icosahedral capsid formed by 72 capsomers, where a capsomer is an association of a pentamer of the VP1 protein linked internally to a single copy of either the VP2 or the VP3 proteins. BKPyV is known to interact with the urothelium and kidney epithelium through the gangliosides GT1b and GD1b but also via other b-series gangliosides (Shinohara et al., 1993; Low et al., 2006; Neu et al., 2013), which are glycosphingolipids carrying one or multiple sialic acids. The crystal structure of a BKPyV VP1 pentamer in interaction with GD3, shows that multiple loops at the surface of the VP1 form a pocket that directly interacts with the α2,8-disialic acid motif of the b-series gangliosides (Neu et al., 2013).

BKPyV is an opportunistic virus with a prevalence of 80% in the worldwide population (Knowles et al., 2003; Egli et al., 2009). Usually, infections occur asymptomatically during childhood and then lead to latency in kidneys. Active viral replication appears to be suppressed by the host, because in immunosuppressive contexts like solid organ or hematopoietic stem cell transplants, BKPyV can reactivate (Hirsch and Steiger, 2003; Hirsch, 2005). This is a particular issue in the case of kidney transplant, where replication in the engrafted kidney results in the secretion of BKPyV in urine (viruria) and the presence of BKPyV DNA in the blood (DNAemia). These parameters are followed clinically as non-invasive markers to evaluate the state of the infection in the graft. Viruria higher than 10^7^ cp/mL and DNAemia greater than 10^4^ cp/mL are associated with BKPyV-associated nephropathy (BKPyVAN), that must be confirmed through biopsy (Hirsch et al., 2013; Nickeleit et al., 1999). Persistent and uncontrolled BKPyV replication can lead to kidney graft dysfunction and ultimately loss of the graft (Ramos et al., 2002; Drachenberg et al., 2007; Viscount et al., 2007). Usually, the therapeutic strategy is to reestablish the host immune response against BKPyV by modulating the immunosuppressive treatments without endangering the graft (Babel et al, 2011). However, for approximately 25% of BKPyVAN patients, high-level BKPyV replication persists (Nickeleit et al., 2018) despite immunosuppressive treatment modulation, and these are the patients with the highest risk of subsequent graft loss (Nickeleit et al., 2021). Persistent high-level BKPyVAN (Randhawa et al., 2002) or DNAemia is accompanied by the accumulation of mutations in the VP1 capsid protein which cluster around the sialic-acid binding pocket (McIlroy et al., 2020).

These mutations appear to be caused by viral genome editing by host APOBEC3 enzymes and lead to neutralisation escape (Peretti et al., 2018), a model in which host innate immune responses supply the mutations that are then selected by the adaptive response. In previous work, we found that VP1 mutations also modified the infectivity of pseudotyped particles, suggesting that neutralisation escape mutations could also modify BKPyV tropism (McIlroy et al., 2020).

In this study, we focus on four variant forms of the VP1 protein coming from three kidney recipients who experienced persistent BKPyV replication after graft despite immunosuppressive treatment modulation. Viruses sampled sequentially from these patients accumulated multiple mutations in the BC-loop region of the VP1 protein, which is involved in the direct interaction of the virus with sialic acids. The BC-loop can be divided in two parts: BC1 and BC2 loops where each part faces in different directions (Neu et al., 2013). Through both functional assays and structural studies, we investigated how these mutations influence both the tropism and the structure of BKPyV, and were able to reveal the involvement of a sialic-acid independent receptor in BKPyV infection.

## Materials & Methods

### Patients and clinical samples

Patients in the present study were transplanted between 2011 and 2014, and had previously been included in a prospective observational study approved by the local ethics committee and declared to the French Commission Nationale de l’Informatique et des Libertés (CNIL, n°1600141). All patients gave informed consent authorising the use of archived urine samples, blood samples and biopsies for research protocols. Anonymised clinical and biological data for these patients were extracted from the hospital databases. BKPyV VP1 mutations occurring in these patients have been previously described (McIlroy et al. Viruses 2020). PyVAN was documented by immunohistochemical staining with mouse monoclonal anti-SV40 T Antigen (clone PAb416, Sigma Aldrich - Saint-Quentin-Fallavier France) diluted 1: 50 with polymer-based EnVision FLEX detection system (Dako K8021, les Ulis - France) utilizing onboard Dako Omnisautomate OMNIS (Dako, Les Ulis - France).

### Protein expression and purification

VP1 pentamer were produced using plasmid pET15b expression vector encoding BKPyV VP1 mutant amino acid sequences from positions 30-300 with an N-terminal hexahistidine tag (His-tag) and a thrombin cleavage site (BioCat Gmbh). The protein was expressed in E.coli BL21 DE3 by IPTG induction. Proteins were purified first by nickel affinity chromatography using a 5 mL HisTrap FF crude column (Cytiva). After protein sample loading, the column was washed with 20 mM Tris pH 7.5, 250 mM NaCl, 10 mM imidazole and 10% glycerol, and proteins were eluted by applying a gradient of elution buffer composed of 20 mM Tris pH 7.5, 250 mM NaCl, 500 mM imidazole and 10% glycerol. For glycan array analysis, the His-tag was retained while for crystallisation this tag was cleaved with 10 U/mg of thrombin protease (Cytiva) for 24h at 20°C with agitation. Cleaved and uncleaved pentamers were finally purified by size-exclusion chromatography on a Superdex 200 16/600 column (Cytiva) by eluting protein with 20 mM HEPES pH 7.5 and 150 mM NaCl.

### Crystallisation and structure determination

BKPyV VP1 pentamers were concentrated to 3-4 mg/mL and crystallised at 20°C by hanging drop vapour diffusion against reservoir solutions of 10-18 % PEG 3.350, 0.1M HEPES pH 7.5, 0.1-0.3 M LiCl (drop size 1 μL of protein/1 μL of reservoir). Crystals were harvested and cryoprotected in reservoir solution supplemented with 20 to 25 % of glycerol for several seconds before flash-freezing them in liquid nitrogen.

Diffraction data were collected at the PXIII beamline of the Swiss Light Source of the Paul Scherrer Institut (Villigen, CH) and processed with XDS (Kabsch, 2010). Structures were solved using molecular replacement with Phaser (CCP4) (Winn et al., 2011) using the wild-type (WT) BKPyV VP1 structure (PDB: 4MJ1) as a model. Refinement was done using Phenix (Liebschner et al., 2019) and the model was built and adjusted in Coot (Emsley et al., 2010).

### Cell culture

HEK 293TT cells, purchased from the National Cancer Institute’s Developmental Therapeutics Program (Frederick, Maryland, USA), were grown in complete DMEM (ThermoFisher) containing 10% FBS (Gibco), 100 U/mL penicillin, 100 μg/mL streptomycin (Dutscher), 1x Glutamax-I (ThermoFisher) and 250 μg/mL Hygromycin B (Sigma). RS cells (Evercyte, Vienna, Austria) were grown in Optipro (ThermoFisher) containing 100 U/mL penicillin, 100 μg/mL streptomycin (Dutscher), 1x Glutamax-I (ThermoFisher). GM95 cells purchased from the RIKEN BRC cell bank were cultured in DMEM with 10% FBS (Gibco), 100 U/mL penicillin, 100 μg/mL streptomycin, and 1x Glutamax-I (ThermoFisher), and LNCaP cells were grown in RPMI medium supplemented with 10% of FBS (Gibco) 100 U/mL penicillin, 100 μg/mL streptomycin (Dutscher), and 1x Glutamax-I (ThermoFisher). Cells were maintained at 37°C in a humidified 5% CO_2_ incubator, and passaged at confluence by trypsinization for 10 minutes with 1x TrypLE Express (ThermoFisher).

### VLP production

BKPyV virus-like particles (VLPs) were prepared following the protocols developed by the Buck lab with slight modifications (Pastrana et al., 2012). Briefly, 1.10^7^ HEK 293TT cells were seeded in a 75 cm^2^ flask in DMEM 10% FBS without antibiotics, then transfected using Lipofectamine 2000 reagent (ThermoFisher) according to manufacturer’s instructions. A total of 36 μg VP1 plasmid DNA was mixed with 1.5 mL of Opti-MEM I (ThermoFisher). 72 μL of Lipofectamine 2000 was diluted in 1.5 mL of Opti-MEM I and incubated for 5 min at room temperature prior to mixing with the diluted plasmid DNA. After 20 min at room temperature, 3 mL of DNA-Lipofectamine complexes were added to each flask containing pre-prepared 293TT cells.

Cells were harvested 48h post transfection by trypsinization and washed once in PBS then resuspended in one pellet volume (“v” μL) of PBS, then mixed with 0.4v μL of 25 U/mL type V Neuraminidase (Sigma). After 15 min at 37°C, 0.125v μL of 10% Triton X-100 (Sigma) was added to lyse cells for 15 min at 37°C. The pH of the lysate was adjusted by addition of 0.075v μL of 1M ammonium sulphate, or sodium bicarbonate if VLPs were to be fluorescence-labelled before ultracentrifugation, then 1 μL of 250 U/μL Pierce Nuclease (Pierce) was added to degrade free DNA. After 3h at 37°C, lysates were adjusted to 0.8M NaCl, incubated on ice for 10 min and centrifuged at 5000*g* for 5 min at 4°C. Supernatant was transferred to a new tube and pellet was resuspended in 2 pellet volumes of PBS 0.8M NaCl, then centrifuged. The second supernatant was combined with the first, then pooled supernatant was re-clarified by centrifuging. Cleared lysate from a T75 flask was labelled with 50 μg Alexa Fluor 647 succinimidyl ester (ThermoFisher Ref A20006) for 1 hour at room temperature. Labelled or unlabeled lysate was layered onto an Optiprep 27%/33%/39% gradient (Sigma) prepared in DPBS/0.8M NaCl, then centrifuged at 175 000 *g* at 4°C overnight in a Sw55TI rotor (Beckman). For labelled VLPs, visible bands were harvested directly, while for unlabelled VLPs, tubes were punctured with a 25G syringe needle, and ten fractions of each gradient were collected into 1.5 mL microcentrifuge tubes. 6.5 μL of each fraction was kept for SDS-PAGE to verify VP1 purity and determine peak fractions for pooling, then PBS 5% bovine serum albumin (BSA) was added to each fraction to a final concentration of 0.1% BSA as a stabilising agent. Peak VP1 fractions were pooled, then the VP1 concentration of each VLP stock was quantified by migrating 5μL on SDS-PAGE, then quantifying the VP1 band by densitometry using a standard curve constructed from a series of 5-fold dilutions of BSA starting at 5 μg/well.

### Pseudovirus production

BKPyV and HPV16 pseudovirus (PSV) particles were prepared following the protocols developed by the Buck lab with slight modifications (Pastrana et al., 2012). Briefly, cell preparation and transfection were performed similarly to BKPyV VLP production. However, instead of transfecting only VP1 plasmid, a total of 36 μg plasmid DNA consisting of 16 μg VP1 plasmid, 4 μg ph2b, 8 μg ph3b and 8 μg pEGFP-N1 was transfected into 293TT cells. 48h after transfection, producer cells were collected by trypsinization. The pellet was washed once in cold PBS then resuspended in 800 μL hypotonic lysis buffer containing 25 mM Sodium Citrate pH 6.0, 1 mM CaCl_2_, 1 mM MgCl_2_ and 5mM KCl. Cells were subjected to sonication in a Bioruptor Plus device (Diagenode) for 10 minutes at 4°C with 5 cycles of 1 min ON / 1 min OFF. Type V neuraminidase (Sigma) was added to a final concentration of 1 U/mL and incubated for 30 min at 37°C. 100 μL of 1M HEPES buffer pH 7.4 (ThermoFisher) was added to neutralise the pH, then 1 μL of 250 U/μL Pierce Nuclease (Pierce) was added before incubation for 2 hours at 37°C. The lysate was clarified by centrifuging twice at 5000 *g* for 5 min at 4°C and PSV was purified in an Optiprep gradient as described for VLP production. After ultracentrifugation and fraction collection, 8 μL of each fraction was removed for qPCR and the peak fractions were pooled, aliquoted and stored at −80°C for use in neutralisation assays.

For quantification of pEGFP-N1 plasmid, 5μL of each fraction was mixed with 5 μL of proteinase K (stock of 2 mg/mL) and 40 μL of sterile water. This solution was incubated at 55°C for 60 min followed by 95°C for 10 min. Then, 1 μL of the solution was used for qPCR using Applied Biosystems 2x SYBR Green Mix (Applied Biosystems). Primers were CMV-F 5’-CGC AAA TGG GCG GTA GGC GTG-3’ and pEGFP-N1-R 5’-GTC CAG CTC GAC CAG GAT G-3’. Thermal cycling was initiated with a first denaturation step at 95°C for 10 min, followed by 35 cycles of 95°C for 15 sec and 55°C for 40 sec. Standard curves were constructed using serial dilutions from 10^7^ to 10^2^ copies of the pEGFP-N1 plasmid per tube.

AAV2-GFP vector was prepared by the Nantes Université CPV core facility (https://sfrsante.univ-nantes.fr/en/technological-facilities/biotherapies/cpv-core-facility).

### Ganglioside supplementation

LNCaP cells were seeded at 4.10^5^ cells/well in a 12-well Falcon plate for binding experiments and at 10^4^ cells/well in 96-well Falcon plate for infectivity experiments.

Gangliosides GM1, GT1b, GD1b, GD1a, GT1a and GD3 (Matreya) were dissolved in chloroform/methanol/water (2:1:0.1) at 1 mg/mL and stored at −20°C. For supplementation, gangliosides in their storage solution were diluted to desired concentration in RPMI containing 20 mM HEPES, 100 U/mL of penicillin, 100 μg/mL of streptomycin and 1x Glutamax-I. Then, ganglioside solutions were sonicated 4*30s and put in open Eppendorf tubes for 3h at 37°C to allow evaporation of chloroform and methanol. Ganglioside solution was then added to cells at a final concentration of 5μM and incubated at 37°C for 18h. Culture medium contained 1% FBS during supplementation.

For VLP binding, after ganglioside incorporation, cells were detached with trypsin/EDTA for 15 min at 37°C. Cells were seeded at 25 μL per well in a 96-well V-bottom plate and incubated for 30 min at 4°C with the different BKPyV VLPs coupled with Alexa Fluor 647. After staining, cells were washed with PBS containing 0.5% FBS by centrifuging at 2 000 rpm for 1 min, 3 times. Fluorescence was measured with a Canto II flow cytometer (Becton Dickinson).

For infectivity assays, after ganglioside supplementation, cells were washed twice with complete medium, then PSVs were inoculated. Infection was observed after 5 days by measuring GFP fluorescence with a Cellomics ArrayScan HCS reader (Thermo Scientific). The percentage of GFP^+^ cells was calculated using Cellomics Scan software with identical settings for all wells analysed in a single experiment. For visualisation, GFP and Hoechst fluorescence contrast was enhanced across all wells in a plate in Cellomics View software, then images of representative fields exported as .png files.

### Enzymatic removal of sialic acid and glycosaminoglycans

293TT cells were seeded at 4.10^5^ cells/well in a 12-well Falcon plate for binding experiments and at 10^4^ cells/well in 96-well Falcon plate for infectivity experiments. 293TT cells were treated for 1h at 37°C, with 0.5U/mL of Neuraminidase V from *Clostridium perfringens* (Sigma N2876) in DMEM + 20 mM HEPES + 0.1% BSA or without Neuraminidase V for control conditions. The VLP binding protocol was the same as previously described. For infectivity, cells were inoculated with different PSVs for 3 hours and then washed 3 times with complete medium before incubation for 72-96 hours at 37°C.

For glycosaminoglycan (GAG) removal, heparinase I/III (H2519-H8891, Merck) or chondroitinase ABC (C2905, Merck) were applied to 293TT cells for 2h in digestion medium (20 mM HEPES, pH 7.5, 150 mM NaCl, 4 mM CaCl2 and 0.1% BSA) at 37°C. Cells were then used for infectivity assays as described for neuraminidase infectivity experiments.

### Inhibition assays with GAGs

293TT cells were incubated with 100 μg/mL of heparin (H4784, Merck) or chondroitin sulphate A/C (C4384, Merk), followed by inoculation with PSVs. Infection was observed after 48h by measuring GFP fluorescence with Cellomics ArrayScan HCS reader (Micropicell, SFR Bonamy). HPV 16 and AVV 2 PSVs carrying eGFP plasmid were used as positive controls.

### Glycan array screening

The binding specificities of the his-tagged recombinant BKPyV VP1s were analysed in the neoglycolipid (NGL)-based microarray system (Liu et al., 2012). Two versions of microarrays were used: (1) ganglioside-focused arrays featuring 26 glycolipid and NGL probes (Fig 2A-C), and (2) broad spectrum screening microarrays of 672 sequence-defined lipid-linked glycan probes, of mammalian and non-mammalian type essentially as previously described (McAllister et al., 2020). Theglycan probes included in the screening arrays and their sequences are given in Supplemental Table 2. Details of the preparation of the glycan probes and the generation of the microarrays are in Supplementary Glycan Microarray Document (Supplemental Table 3) in accordance with the MIRAGE (Minimum Information Required for A Glycomics Experiment) guidelines for reporting of glycan microarray-based data (Liu et al., 2017). The microarray analyses were performed essentially as described (Khan et al., 2014; Neu et al., 2013). In brief, after blocking the slides for 1h with HBS buffer (10 mM HEPES, pH 7.4, 150 mM NaCl) containing 0.33% (w/v) blocker Casein (Pierce), 0.3% (w/v) BSA (Sigma) and 5 mM CaCl2, the microarrays were overlaid with the VP1 proteins for 90 minutes as protein-antibody complexes that were prepared by preincubating VP1 with mouse monoclonal anti-polyhistidine and biotinylated anti-mouse IgG antibodies (both from Sigma) at a ratio of 4:2:1 (by weight) and diluted in the blocking solution to provide a final VP1 concentration of 150 μg/mL. Binding was detected with Alexa Fluor-647-labelled streptavidin (Molecular Probes) at 1 μg/mL for 30 minutes. Unless otherwise specified, all steps were carried out at ambient temperature. Imaging and data analysis are described in the Supplementary MIRAGE document (Supplemental Table 3).

### Glycolipid extraction and purification

HEK-293-TT and RS cells were detached from T75 flasks and washed twice with PBS. 5.10^6^ cells were lyophilized and extracted twice with CHCl_3_/CH_3_OH (2:1, v/v) and once with CHCl_3_/CH_3_OH (1:2, v/v) using intermediary centrifugations at 2500g for 20 min. The combined supernatants were dried under a nitrogen stream, subjected to mild saponification in 0.1 M NaOH in CHCl_3_/CH_3_OH (1:1, v/v) at 37 °C for 2 h and evaporated to dryness. The samples were reconstituted in CH_3_OH/0.1% TFA in water (1:1, v/v) and applied to a reverse phase C18 cartridge (Waters, Milford, MA, USA) on a Interchim^®^ SPE 6.25ws WorkStation. Reverse phase cartridge was equilibrated in the same solvent. After washing with CH_3_OH/0.1% TFA in water (1:1, v/v), GSL were eluted with CH_3_OH, CHCl_3_/CH_3_OH (1:1, v/v) and CHCl_3_/CH_3_OH (2:1, v/v). The elution fraction was dried under a nitrogen stream prior to structural analysis.

### Mass spectrometry analysis of GSL

Isolated glycosphingolipids were permethylated by the sodium hydroxide/DMSO slurry methods (Khoo and Yu, 2010) at room temperature for 2h under agitation. The derivatization was stopped by addition of water and the permethylated glycans were extracted in CHCl3 and washed at least seven times with water. Permethylated glycosphingolipids were analysed by a MALDI-QIT-TOF Shimadzu AXIMA Resonance mass spectrometer (Shimadzu Europe, Manchester, UK) in the positive mode. Samples were prepared by mixing directly on the target 1 μL of glycosphingolipid sample, solubilized in CHCl_3_, with superDHB matrix solution (10 mg/mL dissolved in CHCl_3_/CH_3_OH (1:1, v/v)) and spotted on the MALDI target. The “mid mode” for a mass range of m/z 1000–3000 was used for scanning and laser power was set to 100, with 2 shots at each of the 200 locations within the circular sample spot.

### Statistics

Significant differences seen in the different experiments were calculated through one-way ANOVA followed by Dunnet’s test where one condition per experiment was used as a control group; **** p<0.0001, *** p<0.001, **p<0.01, *p<0.05. All tests were performed using GraphPad Prism 8.

## Results

### VP1 variants show differing infectious profiles compared to the wild-type strain

To investigate the effect of VP1 mutations on BKPyV tropism, we selected BC-loop mutations previously identified in three patients from the Nantes University Hospital KTx cohort with persistent DNAemia caused by subtype Ib2 BKPyV (Supplemental Figure 1). Pseudoviruses carrying the following VP1 variant proteins were generated: the double mutant K69N E82Q (N-Q) from patient 3.4; the E73Q mutant and the triple mutant A72V E73Q E82Q (VQQ) from patient 3.5; and the E73A mutant observed in patient 3.9. Cell lines 293TT and RS were used to test the infectivity of all variant pseudoviruses as well as wild-type subtype Ib2 pseudovirus. These cell lines were chosen for their kidney origin and their expression of SV40 TAg for amplification of reporter gene expression. In 293TT cells, the E73Q variant was as infectious as the WT while E73A and N-Q variants showed 3-fold lower infectivity and VQQ variant was barely infectious (Fig 1A). In RS cells, the VQQ variant had equivalent, or slightly higher infectivity than WT, depending on the experiment, while infectivity of the E73A variant was slightly lower than WT. The strongest differences were observed for the N-Q variant, which was almost non-infectious in this cell line, and the E73Q variant, which showed five to ten-fold higher infectivity compared to the WT (Fig 1B).

**Figure 1.**
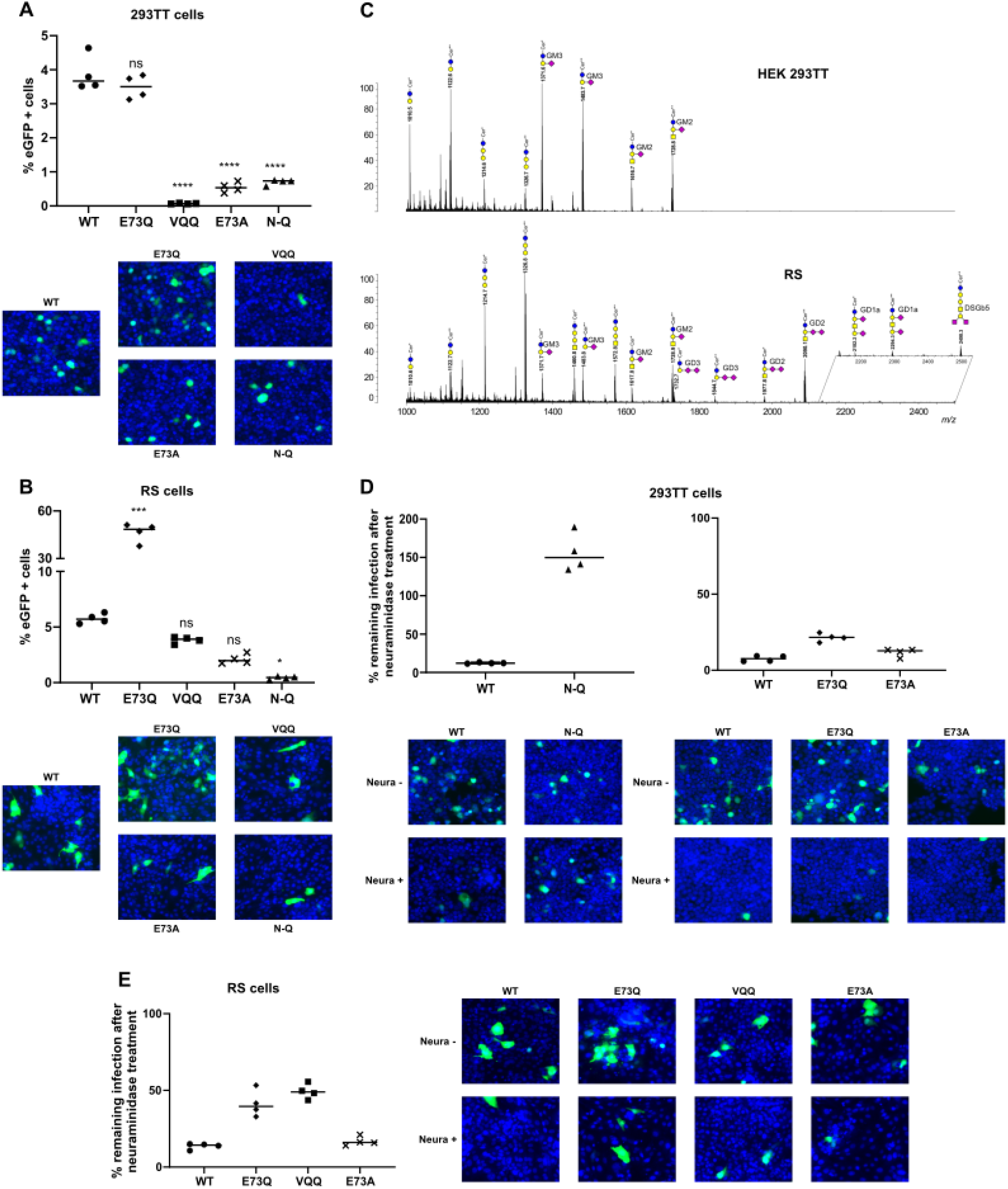
Patient-derived variant of BKPyV have distinct infectious profiles. (A), (B) Infectivity assays in 293TT and RS cells. Cells were inoculated with different variants or WT PSVs. Infection was characterised by quantification of GFP^+^ cells. Significant differences were tested through one-way ANOVA followed by Dunnett’s test with “WT” condition used as the control group **** p<0.0001, *** p<0.001, **p<0.01, *p<0.05. Representative images of cells infected by each PSV in both cell lines are associated with each panel. (C) Comparison of MALDI-TOF-MS profiles of permethylated glycosphingolipids from HEK-293-TT and RS cells. GSL are present as d18:1/C16:0 (Cer*) and d18:1/C24:0 (Cer**) isomers. yellow circles, galactose; blue circles, glucose; yellow squares, *N*-acetylgalactosamine; purple diamonds, *N*-acetylneuraminic acid. (D), (E) Percentage of remaining infection by PSVs after neuraminidase treatment in 293TT and RS cells. Representative images of cells with and without neuraminidase infected with different PSVs are associated with each panel.

To gain insights into the basis of the different infectious profiles that we observed of WT and variant pseudoviruses we characterised the ganglioside profiles of the 293TT and RS cell lines by mass spectroscopy following organic extraction of cell membrane components. Analysis was performed by a combination of MALDI-QIT-TOF-MS and MS/MS of permethylated glycosphingolipids to establish the cellular profiles and the sequence of individual components. Both cell lines were shown to contain monosialylated GM2 and GM3 a-series gangliosides along with neutral globosides. In addition, RS cells, but not 293TT cells, specifically expressed b-series disialylated gangliosides GD2 and GD3 carrying Sia-Sia epitopes as well as lower proportions of the a-series ganglioside GD1a substituted by a single sialic acid on both internal and external Gal residues (Fig 1C). Finally, RS cells exhibited the uncommon disialylated Globo-series ganglioside DSGb5.

From these observations and knowing that the mutations are found in the BC-loop, which is directly involved in ganglioside binding, we sought to characterise their effects on glycan binding specificity.

### N-Q variant infection is sialic acid-independent

Since polyomaviruses are known to interact with sialylated glycans, we tested whether the infectivity of mutant pseudoviruses was also sialic-acid dependent by infecting cells treated with type-V neuraminidase, which removes terminal α-2,3-α-2,6- and α-2,8-linked sialic acid residues (Fig 1D & E). VQQ variant infection was only performed in RS cells while N-Q variant only in 293TT because of the weak infectious capacity of each variant for the other cell line. However, E73Q and E73A variant infectivity was tested in both cell lines. In RS cells, WT, E73Q, VQQ and E73A BKPyV infectivity was significantly decreased by the removal of sialic acid, leading to a remaining infection of around 15% for WT and E73A PSVs, 50% for VQQ and 40% for E73Q PSVs. In 293TT cells, the ability of WT BKPyV to infect was also impacted by the lack of sialic acid on the cell surface with only around 10% of remaining infection for WT and E73A, and around 20% for the E73Q variant. However, the N-Q variant maintained its ability to infect 293TT cells after sialic acid removal, indicating that its infection occurs in a sialic acid-independent manner. Indeed, its infectious ability was even enhanced by sialic acid removal. Thus, another receptor must be involved to support N-Q variant infection in 293TT cells.

### Variants have distinct glycan binding profiles compared to WT

BKPyV is known to use the gangliosides GD1b and GT1b as entry receptors (Low et al, 2006) but is also able to infect cells through other b-series gangliosides (Neu et al, 2013). Gangliosides of the b-series are characterised by their two sialic acids (with an α2-8-linkage) attached to the first galactose of the carbohydrate chain via α2-3-sialyl linkage as in GD3 and GD1b. This disialyl moiety interacts directly with the BKPyV capsid. To assess the binding profile of the BKPyV variants, purified variant and WT genotype I VP1 pentamers were used for glycan array screening with aganglioside-focused array comprising 26 ganglioside-related probes, glycolipids and ganglio-oligosaccharide NGLs (Fig 2A-C). The microarray analysis revealed different binding profiles for theVP1 variants compared to WT VP1 pentamers. The WT BKPyV binding was consistent with previously published data where signal was detected for GT1b and GD1b probes (Neu et al, 2013). A broader profile was observed for the E73A, E73Q, and VQQ variants, comprising binding signals for GD1a, GT1a, and GQ1b, as well as NGL probes GM1b-DH and GD1a-DH in addition to GT1b and GD1b. The structural difference for a-series compared to b-series gangliosides is that they carry a single sialic acid α2-3-linked to the on the first galactose from the reducing end of the carbohydrate chain (Fig 2C). These data indicate that mutations at the E73 residue of the VP1 protein lead to less specific and broader binding, including gangliosides of both band a-series. The overall binding intensities for the two variants with E to Q mutation at position 73, E73Q and VQQ, were higher compared to the E73A and the WT VP1s. For the double N-Q mutant, negligible or no significant binding was observed to any of the probes included in the glycan array indicating the loss of ability to bind to sialylated glycan moities of gangliosides for the N-Q mutant.

**Figure 2.**
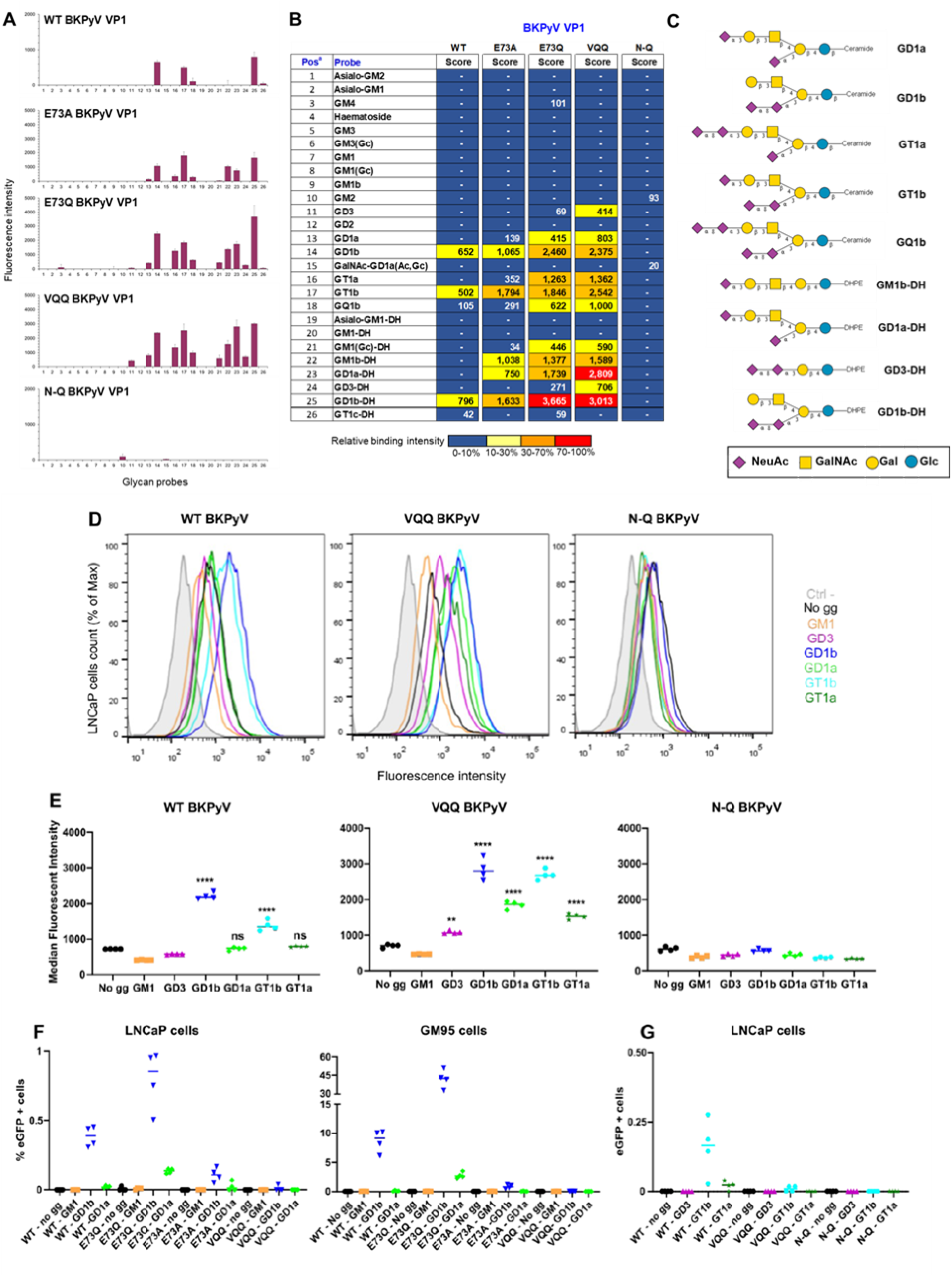
BKPyV variants have distinct glycan binding profiles. (A) Ganglioside-focused glycan microarray results shown as histogram charts of showing fluorescence intensities of binding of His-tagged BKPyV VP1s as means of duplicate spots at 5 fmol/spot. Error bars represent half of the difference between the two values. (B) Heat maps of relative intensities and fluorescence binding scores of BKPyV VP1s (means of duplicate spots at 5 fmol/spot); 100%, the highest binding intensity in the five experiments. (C) Sequences of oligosaccharide probes variously bound by BKPyV VP1s given as symbolic representations. Symbols used for individual monosaccharides are indicated in the key at the bottom of the panel. The full list of the 26 probes with sequences is included in Supplemental Table 3. (D) Histograms representing fluorescent intensities of Alexa-647 variant or WT VLPs during cell-binding assays on ganglioside supplemented LNCaP cells. LNCaPs cells alone are shown in a filled gray histogram for negative control, other conditions with VLPs are represented by coloured histograms. (E) Median fluorescence intensity comparison for binding of each VLP to LNCaP cells supplemented with different gangliosides. Significant differences were tested through one-way ANOVA followed by Dunnett’s test with “no gg” condition used as the control group **** p<0.0001, *** p<0.001, **p<0.01, *p<0.05. (F) (G) Infectivity assays on ganglioside supplemented LNCaP or GM95 cells with different PSVs. Infection was characterised through quantification of GFP^+^ cells.

The five VP1 proteins were further analysed in a broad-spectrum glycan screening array encompassing 672 sequence-defined lipid-linked oligosaccharide probes, representing the major types of mammalian glycans found on glycoproteins (N-linked and O-linked), glycolipids, and proteoglycans, as well as those derived from polysaccharides of bacteria, fungi, and plant origins (Supplemental Fig 2 and Supplemental Table 2). In overall agreement with findings in the ganglioside-focused arrays, the WT, the single and the triple mutant VP1s (E73A, E73Q, and VQQ) showed binding to sialyl glycans but not to the over 400 neutral and sulfated glycan probes that do not contain sialic acids. With the repertoire of sialyl glycans included in the screening array it is clear that the E73Q and VQQ variants acquired the ability to bind to a broader range of sialyl glycans, beyond ganglioside sequences, with enhanced signal intensities compared to the WT VP1 (Supplemental Table 2). The E73A showed a binding pattern similar to that of the WT VP1 except for the additional binding to the a-series ganglioside GD1a and the NGL probe GM1b-DH observed in the focused array. Binding was not detected with the N-Q double mutant VP1 to any sialylated or non-sialylated glycans included in the screening arrays. It remains possible that carbohydrate ligands for the N-Q mutant exist but were not included in the arrayed probe library.

To corroborate the glycan array results, cellular binding assays were performed on LNCaP cells supplemented with a panel of gangliosides (GM1, GD3, GD1b, GT1b, GD1a and GT1a) with fluorescent WT, VQQ and N-Q VLPs (Fig 2D-E). WT VLPs bound to cells supplemented with GD1b and GT1b, consistent with the known role of these gangliosides as BKPyV receptors. VQQ VLPs were able to bind all gangliosides tested except GM1, confirming the glycan array data. Stronger binding was seen for both GD1b and GT1b followed by GD1a and GT1a and then weak binding to cells supplemented with GD3 (Fig 2D-E). Also consistent with the glycan array results, N-Q VLPs did not bind any of the tested gangliosides (Fig 2D-E).

### Gangliosides are not sufficient to support VQQ variant infection

To determine whether binding of the different gangliosides by WT, VQQ, E73Q and E73A variant VP1 can lead to infection, we performed infectivity assays on LNCaP and GM95 cells supplemented with gangliosides. As expected, infection was seen for WT in cells supplemented with GD1b, but also for E73Q and E73A PSVs under the same condition, and infection was also seen for E73Q when cells were supplemented with GD1a in both LNCaP and GM95 cells (Fig 2F). The ability of the E73Q variant to infect cells supplemented with GD1a was consistent with its increased infectivity in RS cells, which carried GD1a gangliosides. Surprisingly, no infection was observed for VQQ in any supplementation condition, despite its ability to bind GD1b, GD1a, and other gangliosides on the surface of cells. Hence, ganglioside binding was not sufficient for infection with VQQ variant PSV. Finally, as predicted, no infection was seen in any supplementation condition for the N-Q variant, confirming that gangliosides are not used by this variant for infectious entry (Fig 2G).

### Mutations at VP1 positions 72 and 73 can lead to BC-loop flipping

To understand the impact of the mutations on the VP1 protein structure, variant structures were solved through X-ray crystallography. The single mutants E73Q, E73A, and triple mutant VQQ VP1 pentamer structures were solved at 1.85, 1.89 and 1.8 Å (Table 1). All variant VP1 pentamer structures were globally similar to the WT VP1 pentamer with a doughnut-shaped ring composed of a central pore surrounded by five VP1 monomers in five-fold symmetry. Like the WT structure, the monomers have a β-sandwich fold with jelly-roll topology. The eight β-strands (B, I, D, G and C, H, E, F) are linked by extensive loops, exposed at the surface of the protein (Fig 3A-B).

**Figure 3.**
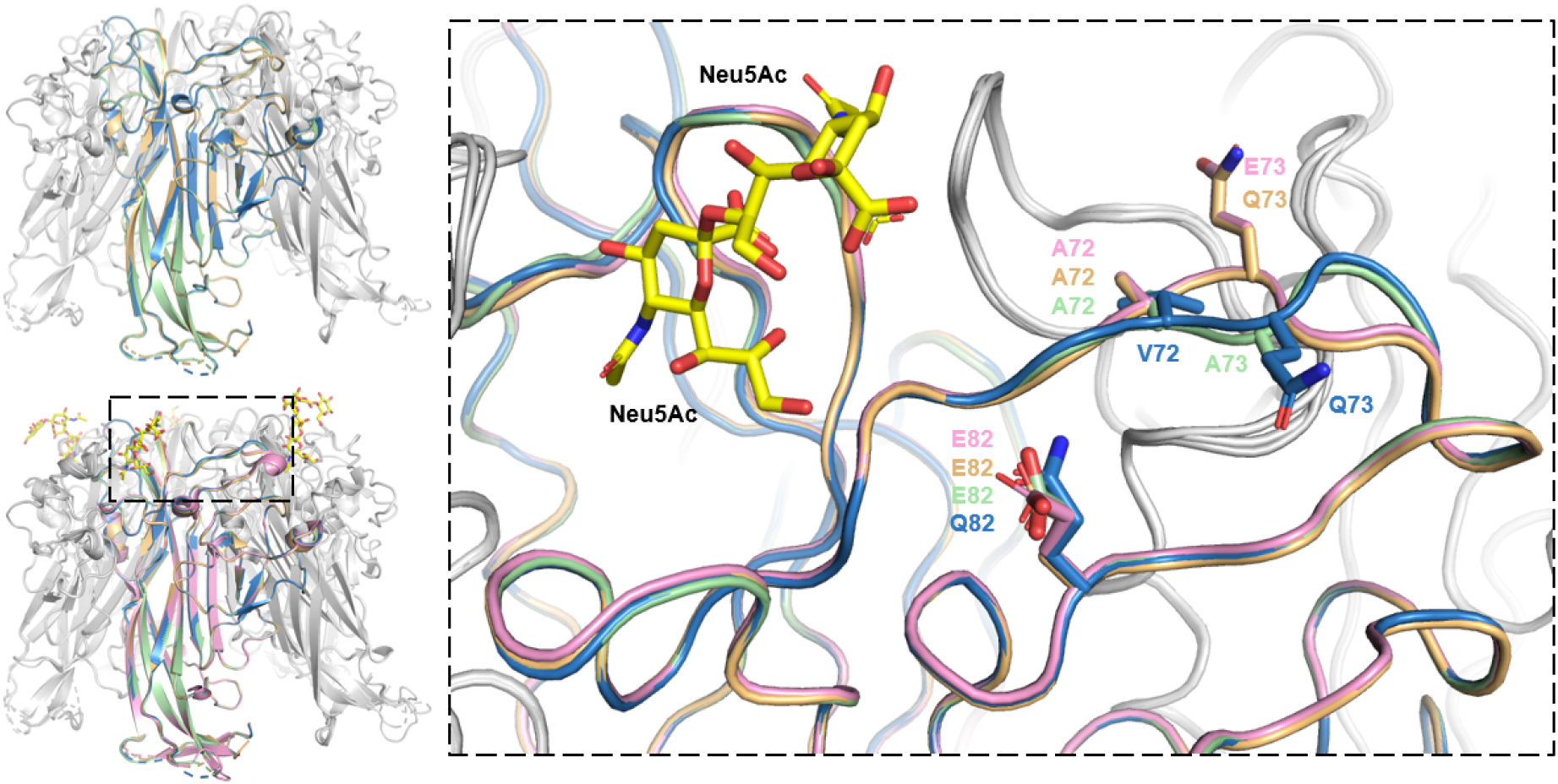
Mutations of amino acids 72 and 73 lead to structural changes in the BC-loop conformation. (A) Superposition of variant E73Q, E73A and VQQ VP1 pentamer structures. One monomer for each structure is highlighted in yellow for E73Q, green for E73A and blue for VQQ. (B) Superposition to variant VP1 pentamers with BKPyV WT VP1 pentamer in association with GD3 oligosaccharides. WT VP1 monomer is highlighted in pink and GD3 molecules are represented by yellow sticks. (C) Zoom on the BC-loops of highlighted VP1 monomers. Amino acids 72, 73 and 82 are represented as sticks.

**Table 1.** Data processing and refinement.

| **Data collection** | **gl E73Q** | **gl E73A** | **gl A72V E73Q E82Q** | **gl K69N E82Q** |
| --- | --- | --- | --- | --- |
| Beamline |  |  | PXIII |  |
| Wavelength (Å) |  |  | 1 |  |
| Space group | P2(1)2(1)2 | P2(1)2(1)2 | P2(1)2(1)2 | P2(1) |
|  | a = 145.25 | a = 144.49 | a = 144.69 | a = 60.46 |
| Unit cell length (Å) | b = 153.04 | b = 152.30 | b = 152.60 | b = 155.96 |
|  | c = 62.80 | c = 62.58 | c = 62.75 | c = 141.32 |
| | $\alpha = 90$ | $\alpha = 90$ | $\alpha = 90$ | $\alpha = 90$ |
| Unit cell angle (°) | $\beta = 90$ | $\beta = 90$ | $\beta = 90$ | $\beta = 92.64$ |
| | $\gamma = 90$ | $\gamma = 90$ | $\gamma = 90$ | $\gamma = 90$ |
| Molecules/ASU | 1 | 1 | 1 | 2 |
| Resolution(Å) | 50 - 1.85 (1.87-1.85) | 50 - 1.89 (1.91-1.89) | 50 - 1.80 (1.82-1.80) | 50 - 2.01 (2.03-2.01) |
| Completeness (%) | 99.8 (99.5) | 99.9 (99.4) | 99.9 (99.6) | 94.1 (97.1) |
| Total reflections | 1 579 907 (251 766) | 1 494 946 (239 425) | 1 743 939 (284 486) | 1 088 105 (159 494) |
| Unique reflections | 119 355 (19 035) | 111 734 (17 862) | 129 673 (20 735) | 164 929 (27 282) |
| Rmeas (%) | 17.2 (162.6) | 13.2 (144.7) | 14.7 (154.7) | 20.8 (127.9) |
| CC1/2 (%) | 99.9 (75.7) | 99.9 (69.1) | 99.8 (64.8) | 99.5 (99.5) |
| I/$\sigma$ | 14.02 (1.69) | 15.56 (1.96) | 13.40 (1.68) | 8.33 (1.39) |
| **Refinement** |  |  |  |  |
| Rwork/Rfree (%) | 18.86/23.05 | 17.38/21.11 | 17.12/21.00 | 17.98/23.13 |
| RMS deviation: |  |  |  |  |
| Bond length (Å) | 0.010 | 0.010 | 0.011 | 0.010 |
| Bond angle (°) | 1.60 | 1.62 | 1.65 | 1.71 |
| Average B factor (Å<sup>2</sup>) | 31.0 | 33.0 | 29.0 | 31.0 |
| Wilson B factor (Å<sup>2</sup>) | 25.8 | 28.6 | 24.6 | 25.9 |

The E73Q pentamer structure was almost identical to that of the WT (r.m.s.d. of 0.5 Å), whereas the E73A mutant presenting an alanine in the same position, showed some structural changes (r.m.s.d of 1.3 Å). Indeed, a shift of the BC2-loop, which includes the E73A mutation, was clearly seen for two monomers out of the five (Fig 3C). However, two monomers retained the WT loop conformation while the last monomer seemed to have a state where both conformations are superimposed. Thus, this mutation to alanine induces a more flexible conformation for the loop compared to the WT and the E73Q single mutant. Interestingly, while the E73Q mutation on its own did not induce any structural changes, when combined with A72V, as seen in the VQQ variant, the second part of the BC-loop was switched (r.m.s.d of 1.5Å) in a similar manner to that seen in the E73A variant (Fig 3C). However, in the case of the VQQ variant, the modified orientation of the BC-loop was observed in all five VP1 monomers, not just in the two monomers seen in E73A. This suggests that mutations at position 73 on their own may not be enough to induce a clear structural change in the loop orientation, which only occurs if mutations in this position are combined with the A72V mutation at the neighbouring amino acid.

However, structural changes seen in the VQQ variant are not directly involved in the ganglioside interaction. As previously described, the double sialic part of the b-series gangliosides interacts with the BC1-loop, not the BC2-loop, as well as HI- and DE-loops (Neu et al, 2013). Thus, the structure of the VQQ variant is consistent with its ability to bind gangliosides.

### K69N mutation in N-Q variant induces loss of interaction with sialic acids

The structure of the N-Q variant VP1 pentamer was also solved at 2.01 Å (Table 1). Like the other variants, the N-Q VP1 pentamer structure was very similar to the WT VP1 pentamer (Fig 4A-B). Although no major change was observed in the backbone orientation of the BC-loop that carried the K69N and E82Q mutations, the mutation from lysine to asparagine at position 69 induced significant changes (Fig 4C). Lysine 69 is involved directly with the double sialic acid part of the b-series gangliosides by making van der Waals, salt bridge and hydrogen bond interactions through its side chain, and its replacement by asparagine leads to loss of those interactions (Fig 4D). Moreover, if gangliosides were to bind the pentamer in the same way, the asparagine side chain would be orientated too close (1.3 Å) to the hydroxyl group on C8 of the second sialic acid, leading to a steric clash between the sialic acid and the asparagine.

**Figure 4.**
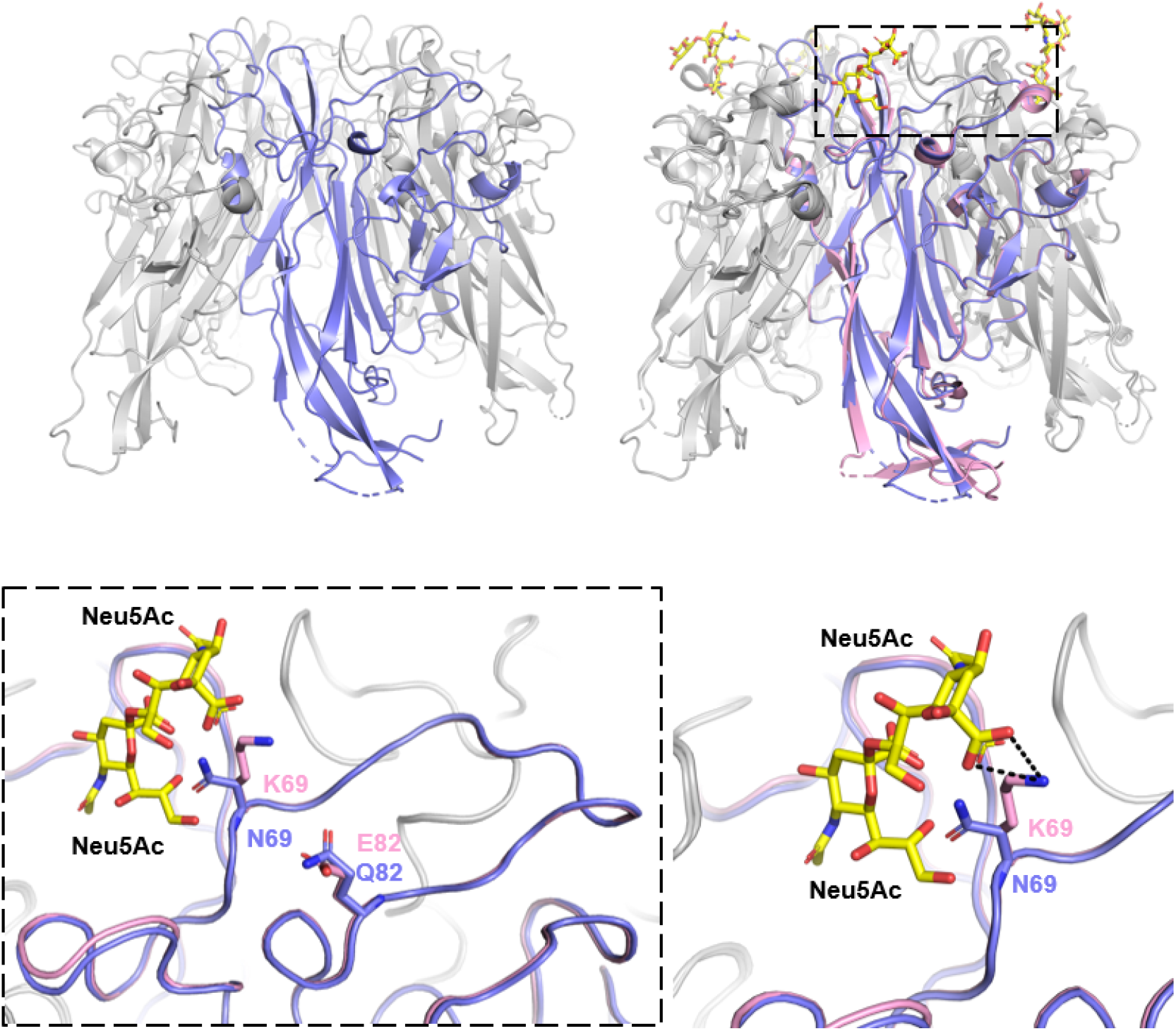
Structure of N-Q VP1 pentamer. (A) Structure of N-Q VP1 pentamer. VP1 monomer is highlighted in purple. (B) Superposition of N-Q VP1 pentamer onto WT VP1 pentamers associated with GD3 oligosaccharides. WT VP1 monomer is highlighted in pink and GD3 molecules are represented by yellow sticks. (C) Zoom on the BC-loops of highlighted VP1 monomers. Amino acids 69 and 82 are represented as sticks. (D) Focus on amino acids in position 69 and their interaction with Neu5Ac. Hydrogen bonds represented by black dashed lines between K69 and hydroxyl groups of Neu5Ac.

### WT and N-Q BKPyV do not interact with GAGs

Despite losing its ability to bind sialic acid, the N-Q variant remains infectious in the 293TT cell line, indicating that another receptor is used by this variant to enter these cells. Merkel Cell Polyomavirus (MCPyV) is known to interact with both sialylated and non-sialylated glycans for infection. Like BKPyV, MCPyV can interact with gangliosides as attachment receptors, but also heparan sulphate (HS) or chondroitin sulphate (CS) glycosaminoglycans (GAGs), as entry receptors (Schowalter et al., 2011). To test whether the BKPyV N-Q variant could interact with GAGs as an alternative entry receptor, heparin and chondroitin sulphate A/C were added to 293TT cells before and during the infection experiments. Pre-incubation with heparin effectively blocked infectious entry of HPV16 and AAV2, which are both known to use GAGs as entry receptors, whereas pre-incubation with chondroitin sulphate significantly reduced infectious entry of HPV16 (Fig 5A). In contrast, neither heparin, nor chondroitin sulphate had any effect on WT or N-Q BKPyV infectivity (Fig 5). To further confirm that GAGs are not receptors for N-Q variant capsids, 293TT cells were treated with heparinase I/III or chondroitinase ABC to remove heparan sulphate or chondroitin sulphate from the cell surface. Infection by both WT and N-Q PSVs was not inhibited by the enzyme treatment (Fig 6B).

**Figure 5.**
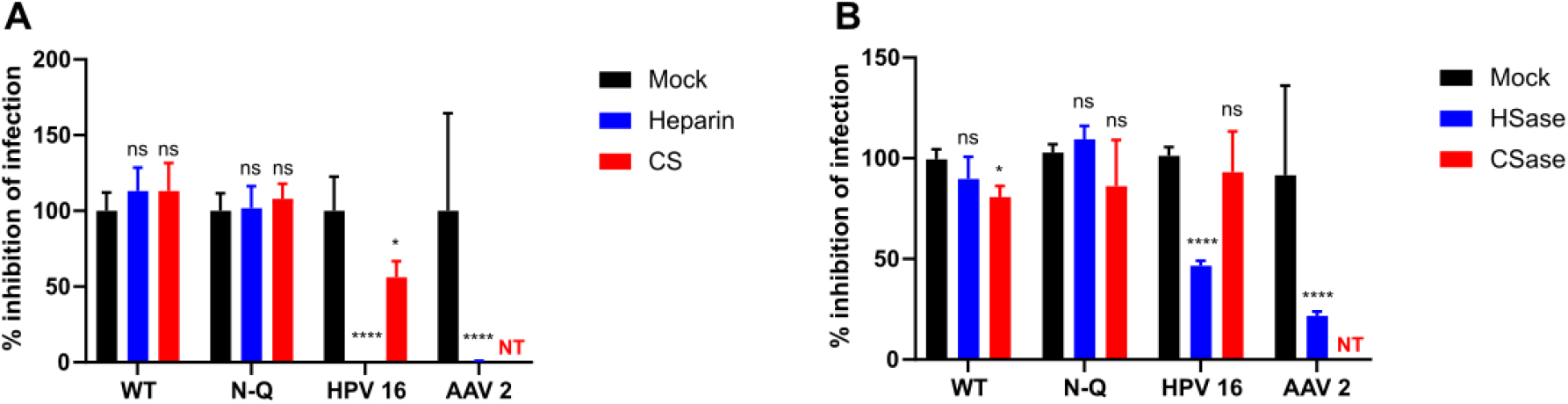
N-Q does not interact with GAGs. (A) Infectivity assays in presence or not of 100 μg/ml of heparin or chondroitin sulfate A/C with BKPyV WT and N-Q PSVs. HPV 16 and AAV 2 PSVs serve as positive control. Error bars correspond to standard deviations. NT - Not tested. (B) Infectivity assays treated or not with heparinase I/III or chondroitinase ABC with BKPyV WT and N-Q PSVs. HPV 16 and AAV 2 PSVs serve as positive control. Error bars correspond to standard deviations. NT - Not tested.

**Figure 6.**
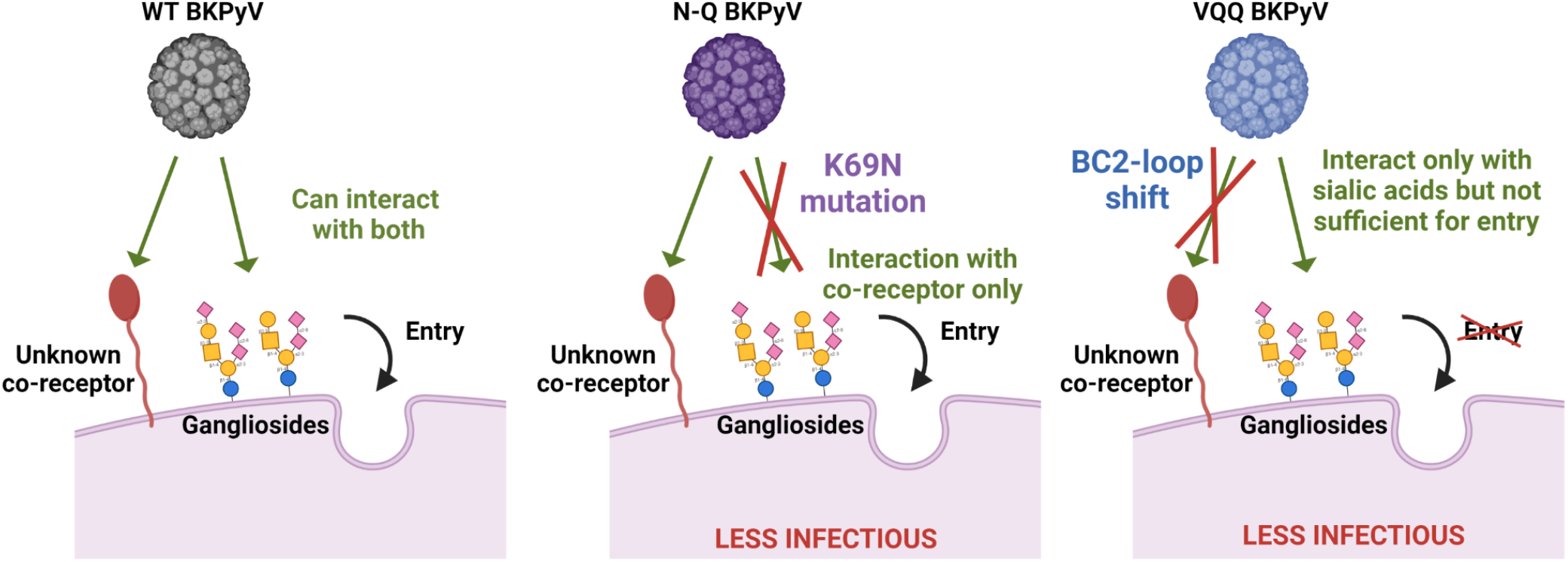
Putative entry mechanisms of N-Q and VQQ variants.

## Discussion

In this functional and structural study, we show that mutations in the BC-loop of the VP1 protein that occur in KTx recipients impact the ganglioside binding specificity of the BKPyV capsid, and we describe two very different functional patterns, illustrated by the VQQ and N-Q variants.

The N-Q variant has lost the ability to interact with sialic acids due to the mutation of lysine 69 to asparagine. In a previous study, mutation of the same lysine to serine induced a change of tropism from b-series gangliosides to GM1, known to be an SV40 receptor (Neu et al., 2013). This emphasises the essential role that amino acid 69 seems to play in the interaction with gangliosides. Pseudotypes carrying N-Q VP1 had reduced infectivity in both RS and 293TT cells, but the remaining infectivity in 293TT cells was entirely sialic-acid independent, and in fact increased after neuraminidase treatment, suggesting that an alternative receptor is involved in infectious entry of the N-Q variant. Our first candidates were the heparan sulphate and chondroitin sulphate GAGs, since it has been reported that MCPyV (Schowalter et al, 2011) and mutant JCPyV and BKPyV pseudotypes can use GAGs for infectious entry (Geoghagan et al. 2017). However, neither the addition of excess heparin or chondroitin sulphate nor the removal of HS or CS GAGs from 293TT cells had any effect on the infectivity of WT or N-Q BKPyV. Furthermore, none of the BKPyV VP1 variants bound to any of the glycosaminoglycan oligosaccharides presented on the broad spectrum glycan array. We therefore tentatively conclude that an as yet uncharacterized receptor is involved in the infectious entry of the N-Q BKPyV variant.

Mutations at VP1 residue 73 had two distinct effects on VP1 structure: first, loss of charge at the capsid surface on the lip of the sialic acid binding site, seen in the E73A, E73Q and VQQ variants; and secondly, a conformational flip of the BC2-loop with the orientation of amino acids 73 and 74 significantly shifted. Mutation of the initial glutamic acid at position 73 to glutamine in the E73Q variant did not modify the orientation of the BC-loop, while its mutation to alanine, in E73A, revealed the flexibility of this part of the BC loop. In the E73A variant crystal structure, two VP1 monomers kept their initial BC-loop conformation, two monomers had an alternative orientation of the loop, and the last monomer appeared to have both conformational states of the BC2-loop superposed. In the crystal structure of the VQQ variant, all five monomers showed the alternative, “flipped” conformation. Thus, we concluded that on its own, mutation of amino acid 73 does not induce a stable conformational change of the loop, which requires a second mutation at position 72. The requirement for the A72V mutation in order to stabilise the alternative orientation of BC-loop region 73-75 explains the previously reported statistical association of the A72V mutation with E73 mutations in patient samples (McIlroy et al., 2020), and is consistent with the sequential appearance of VP1 mutations - first at position 73, then followed by A72V, that were seen in patients 3.5 and 3.9 (Figure S1).

Variants E73A, E73Q and VQQ all showed broader ganglioside binding (Figure 2A), indicating that negative charge at residue 73 may restrict WT BKPyV VP1 binding to b-series gangliosides, and that removing this charge is sufficient to allow binding to a-series gangliosides. On the other hand, only variant E73Q showed strongly enhanced infectivity in RS cells which, as shown by mass spectroscopy, carried the GD1a ganglioside, and only variant E73Q was able to infect LNCaP and GM95 cells supplemented with GD1a. Similarly, in 293TT cells, E73Q retained infectivity equivalent to that of WT VP1, whereas both VQQ and E73A showed significantly reduced infectivity. Therefore, flipping of the BC2-loop, and not loss of negative charge at position 73, was correlated with loss of infectivity in 293TT cells. This led us to a model in which the two different structural effects of VP1 mutations at position 73 have opposite effects on infectivity: loss of charge increases infectivity, by broadening ganglioside binding, whereas BC-loop flipping reduces infectivity. These two effects explain the infectivity patterns that were observed for the different VP1 variants. In RS cells, the ability to use GD1a as an entry receptor results in the strongly increased infectivity of the E73Q variant, whereas for E73A and VQQ, enhanced ganglioside binding was balanced by a loss of infectivity due to BC-loop flipping, leading to near-WT infectivity for both variants. In 293TT cells, although the infectivity of WT, E73Q and E73A variants was sialic acid dependent, this was not due to gangliosides, since neither a- nor b-series gangliosides were present on 293TT cells (Figure 1C). Therefore, the infectivity of E73Q was not increased relative to wild type, and the negative impact of BC-loop flipping on the infectivity of E73A and VQQ variants was not compensated by increased ganglioside binding. One previous report described sialic acid dependent infection by BKPyV that was strongly impacted by tunicamycin, an inhibitor of N-linked protein glycosylation, leading the authors to conclude that an N-linked glycoprotein, rather than a ganglioside, was an entry receptor for BKPyV (Dugan et al., 2005). A receptor of this type could be involved in infectious entry in 293TT cells. In the broad-spectrum glycan screening arrays, indeed there was binding with various intensities to sialyl glycans beyond ganglioside sequences (Supplemental Table 2). These could be followed up further in future structural and functional studies.

Overall, the N-Q and VQQ variants appeared to be almost mirror images of each other. The N-Q variant lost all ganglioside-binding activity, but retained infectivity in 293TT cells through a sialic-acid independent pathway, whereas VQQ showed enhanced ganglioside binding, but almost completely lost infectivity in 293TT cells. One plausible explanation of these observations is that the VQQ variant may have lost the ability to interact with the unknown entry receptor employed by the N-Q variant to infect 293TT cells, and that this interaction is required, in addition to sialic acid binding, for infectious entry (Figure 6). This would explain why binding and infectivity were uncoupled for the VQQ variant in both 293TT cells and ganglioside-supplemented LNCaP and GM95 cells. If this is correct, then the WT orientation of the BC-loop residues 73-74 must be critical for the interaction of the BKPyV capsid with this putative receptor.

What are the mutational and selective pressures that drive the emergence of these mutations in patients? As previously noted, the E73Q and E82Q mutations we analysed occur at optimal APOBEC3A/B editing sites (Peretti et al., 2018), however, although both the A72V and K69N mutations involve cytosine deamination, they are located in the trinucleotide context G<u>C</u>T, which is not an APOBEC3A/B target site. In addition, the E73A mutation involved a T>G transversion. These observations imply that additional mutagenic processes, independent of APOBEC3A/B, occur in KTx recipients. Once they arise, BC-loop mutations conferring neutralisation escape appear to be selected by the host humoral response, (Peretti et al., 2018; McIlroy et al., 2020), which explains why variants with reduced infectivity relative to WT can rise to dominance in a given individual’s viral population. In terms of tissue tropism, although patient 3.4 underwent transplantectomy, due to deteriorating graft function, at 49 months post-transplant, BKPyV-infected cells could not be clearly identified in kidney tissue at this time (Supplemental Figure 1A). In contrast, PyVAN was clearly diagnosed in patient 3.9 at a time when the viral population was predominantly WT (Supplemental Figure 1C), whereas graft function was stable, with no biopsies taken, after the patient’s second graft, when the E73A mutation dominated the viral population. Biopsies were not available for patient 3.5 at time points when the VQQ variant was documented, although the graft was still functional, without PyVAN at 70 months post-graft. These observations are consistent with the reduced *in vitro* infectivity of the VQQ and N-Q variants, and suggest that BKPyV carrying the K69N mutation had an attenuated phenotype *in vivo.*

Our study has some important limitations, in particular the description of the structural and functional impact of BKPyV VP1 mutations remains incomplete because additional VP1 mutations accumulated in patients 3.4, 3.5 and 3.9 over time. In patient 3.5, K69M and D75N were added to the VQQ variant, and in patient 3.4 D60N and A72V were added to N-Q. However, we were unable to express the M^69^V^72^Q^73^N^75^Q^82^ and the N^60^N^69^V^72^Q^82^ mutants as either crystallisable VP1 pentamers, VLPs or high-titer pseudotypes. It was therefore not possible to determine whether subsequent mutations induced more radical changes in virus tropism, or on the contrary, compensated for the deleterious effects on infectivity that we document in the VQQ and N-Q variants. Moreover, we were not able to generate structural data from crystals of the VQQ variant complexed with GD1a ganglioside, so the details of this molecular interaction remain to be determined. Nevertheless, the identification of a naturally occurring “sialic acid blind” VP1 variant does allow us to conclude that a sialic acid independent entry receptor for BKPyV, relevant for *in vivo* infection, must exist.

## Acknowledgements

This work was supported by grants from the Agence Nationale de la Recherche (BKNAB ANR-17-CE17-0003), the Université Franco-Allemande (CT-12-20), the German Research Foundation DFG Research Unit FOR2327 ViroCarb, and the Wellcome Trust Biomedical Resource Grants (WT099197/Z/12/Z, 108430/Z/15/Z and 218304/Z/19/Z). The March of Dimes Prematurity Research Center grant 22-FY18-82 provided partial financial support to the Imperial College Glycosciences Laboratory. The sequence-defined glycan microarrays contain many saccharides provided by collaborators whom we thank as well as Professor Ten Feizi and other members of the Glycosciences Laboratory for their contribution in the establishment of the neoglycolipid-based microarray system. YL is grateful to Professor Barbara Mulloy and for the excellent facilities provided by Tom Frenkiel and his team (UK MRC Biomedical NMR Centre at the Francis Crick Institute) for conducting the NMR analysis of oligosaccharides for the Carbohydrate Microarray Facility over the years. For microscopy, we acknowledge the IBISA MicroPICell facility (Biogenouest), member of the national infrastructure France-Bioimaging supported by the Agence Nationale de la Recherche (ANR-10-INBS-04). TS is grateful for beam time at the Swiss Light Source (Villigen, Switzerland).

## Supplemental Data

**Supplemental Table 1.** Age, sex; drug regimen; deceased/live graft; HLA mismatches; Time at 1st +ve viremia; PyVAN; virus genotype. Type of kidney disease? BKneut tire at M0; BK neut titre M12.

|  |  |  |  |
| --- | --- | --- | --- |
| Patient code | 3.4 | 3.5 | 3.9 |
| Age at KTx | 64 | 27 | 56 |
| Sex | M | M | M |
| Graft type | Kidney 1st graph | Kidney 1st graft | Kidney 2nd graft* |
| Donor | Deceased | Deceased | Living familial donor |
| HLA mismatches | 3 | 4 | 4 |
| Immunosuppressive drug regimen | MMF Tacrolimus | MMF Tacrolimus<br>Corticoids | Azathioprine<br>Tacrolimus |
| Log10 BKPyV gIb2 neutralizing titre at KTx | 2.7 | 3.3 | 4.6 (Before KTx2)<br>Serum at KTx1 not available |
| Log10 BKPyV gIb2 neutralizing titre at M12 post-KTx | 4.1 | 5.1 | 4.4 (After KTx2) |

\*The first graft was kidney and pancreas but the first graft ended with PyVAN.

Supplemental Table 2 - List of glycan probes in the broad spectrum glycan screening arrays, their fluorescence binding scores and relative binding intensities (‘matrix’) elicited with BKPyV VP1s (Excel file)

Supplemental Table 3 - Supplementary glycan microarray document based on MIRAGE Glycan Microarray guidelines (PDF file)

**Supplemental Figure 1.**
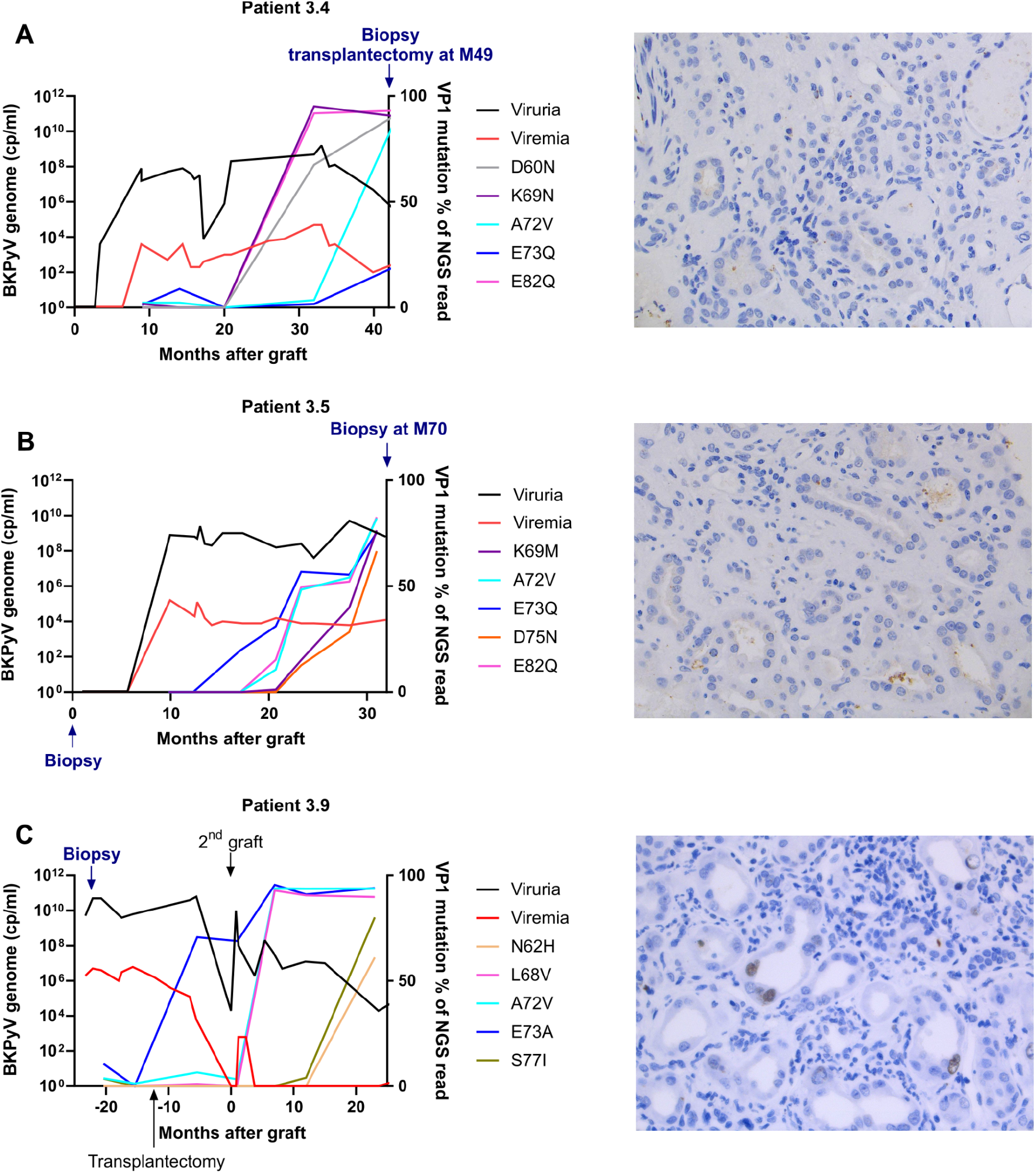
Left panels: Viruria, DNAemia and VP1 mutation % of NGS reads as a function of time after graft for the three kidney recipients whose VP1 variants were studied. (A) Patient 3.4; (B) Patient 3.5; (C) Patient 3.9. Right panels: SV40 TAg staining of biopsies from patients 3.4 - no significant staining; 3.5 - no significant staining; and 3.9 - focal nuclear positivity of epithelial tubular cells.

**Supplemental Figure 2.**
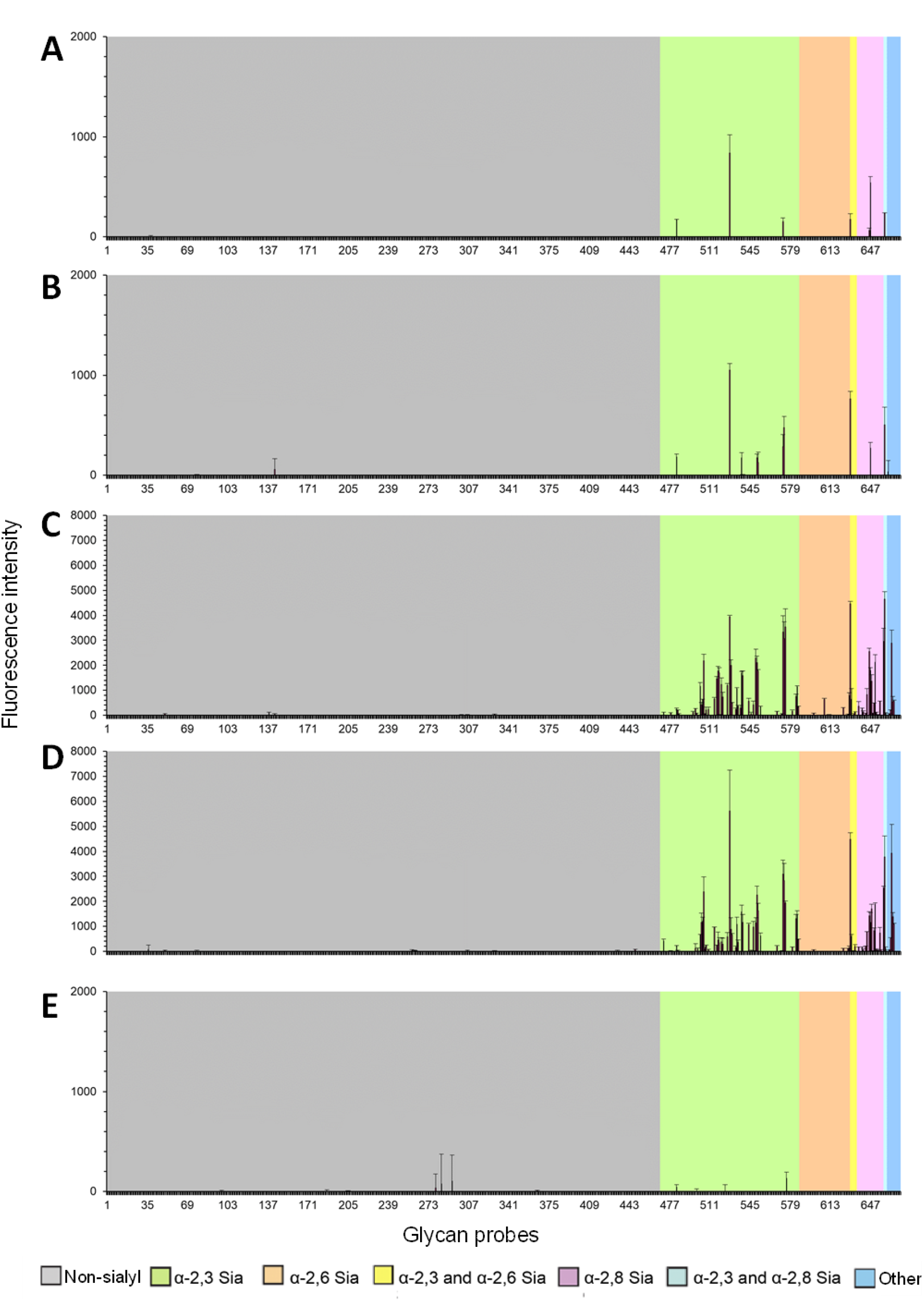
Glycan microarray screening analyses of His-tagged pentameric BKPyV VP1 proteins, wt (A), E73A mutant (B), E73Q mutant (C), VQQ mutant (D) and N-Q mutant (E). The results are the means of fluorescence intensities of duplicate spots, printed at 5 fmol per spot. The error bars represent half of the difference between the two values. In the glycan array the 672 lipid-linked probes are grouped according to sialyl linkages as annotated by the colored panels. The list of glycan probes, their sequences and binding scores are in Supplemental Table 2.

## References

Drachenberg, C.B., Hirsch, H.H., Papadimitriou, J.C., Gosert, R., Wali, R.K., Munivenkatappa, R., Nogueira, J., Cangro, C.B., Haririan, A., Mendley, S., et al. (2007). Polyomavirus BK versus JC replication and nephropathy in renal transplant recipients: a prospective evaluation. Transplantation 84, 323–330. https://doi.org/10.1097/01.tp.0000269706.59977.a5.

Dugan, A.S., Eash, S., and Atwood, W.J. (2005). An N-linked glycoprotein with alpha(2,3)-linked sialic acid is a receptor for BK virus. J. Virol. 79, 14442–14445. https://doi.org/10.1128/jvi.79.22.14442-14445.2005.

Egli, A., Infanti, L., Dumoulin, A., Buser, A., Samaridis, J., Stebler, C., Gosert, R., and Hirsch, H.H. (2009). Prevalence of polyomavirus BK and JC infection and replication in 400 healthy blood donors. J. Infect. Dis. 199, 837–846. https://doi.org/10.1086/597126.

Emsley, P., Lohkamp, B., Scott, W.G., and Cowtan, K. (2010). Features and development of Coot. Acta Crystallogr. D Biol. Crystallogr. 66, 486–501. https://doi.org/10.1107/S0907444910007493.

Hirsch, H.H. (2005). BK virus: opportunity makes a pathogen. Clin. Infect. Dis. Off. Publ. Infect. Dis. Soc. Am. 41, 354–360. https://doi.org/10.1086/431488.

Hirsch, H.H., and Steiger, J. (2003). Polyomavirus BK. Lancet Infect. Dis. 3, 611–623. https://doi.org/10.1016/S1473-3099(03)00770-9.

Hirsch, H.H., Randhawa, P., and the AST Infectious Diseases Community of Practice (2013). BK Polyomavirus in Solid Organ Transplantation: BK Polyomavirus in Solid Organ Transplantation. Am. J. Transplant. 13, 179–188. https://doi.org/10.1111/ajt.12110.

Kabsch, W. (2010). XDS. Acta Crystallogr. D Biol. Crystallogr. 66, 125–132. https://doi.org/10.1107/S0907444909047337.

Khan, Z.M., Liu, Y., Neu, U., Gilbert, M., Ehlers, B., Feizi, T., and Stehle, T. (2014). Crystallographic and Glycan Microarray Analysis of Human Polyomavirus 9 VP1 Identifies N-Glycolyl Neuraminic Acid as a Receptor Candidate. J. Virol. 88, 6100–6111. https://doi.org/10.1128/JVI.03455-13.

Khoo, K.-H., and Yu, S.-Y. (2010). Chapter One - Mass Spectrometric Analysis of Sulfated N-and O-Glycans. In Methods in Enzymology, M. Fukuda, ed. (Academic Press), pp. 3–26.

Knowles, W.A., Pipkin, P., Andrews, N., Vyse, A., Minor, P., Brown, D.W.G., and Miller, E. (2003). Population-based study of antibody to the human polyomaviruses BKV and JCV and the simian polyomavirus SV40. J. Med. Virol. 71, 115–123. https://doi.org/10.1002/jmv.10450.

Liebschner, D., Afonine, P.V., Baker, M.L., Bunkóczi, G., Chen, V.B., Croll, T.I., Hintze, B., Hung, L.W., Jain, S., McCoy, A.J., et al. (2019). Macromolecular structure determination using X-rays, neutrons and electrons: recent developments in Phenix. Acta Crystallogr. Sect. Struct. Biol. 75, 861–877. https://doi.org/10.1107/S2059798319011471.

Liu, Y., Childs, R.A., Palma, A.S., Campanero-Rhodes, M.A., Stoll, M.S., Chai, W., and Feizi, T. (2012). Neoglycolipid-based oligosaccharide microarray system: preparation of NGLs and their noncovalent immobilization on nitrocellulose-coated glass slides for microarray analyses. Methods Mol. Biol. Clifton NJ 808, 117–136. https://doi.org/10.1007/978-1-61779-373-8_8.

Liu, Y, McBride, R., Stoll, M., Palma, A.S., Silva, L., Agravat, S., Aoki-Kinoshita, K.F., Campbell, M.P., Costello, C.E., Dell, A., et al. (2017). The minimum information required for a glycomics experiment (MIRAGE) project: improving the standards for reporting glycan microarray-based data. Glycobiology 27, 280–284. https://doi.org/10.1093/glycob/cww118.

Low, J.A., Magnuson, B., Tsai, B., and Imperiale, M.J. (2006). Identification of Gangliosides GD1b and GT1b as Receptors for BK Virus. J. Virol. 80, 1361–1366. https://doi.org/10.1128/JVI.80.3.1361-1366.2006.

McAllister, N., Liu, Y., Silva, L.M., Lentscher, A.J., Chai, W., Wu, N., Griswold, K.A., Raghunathan, K., Vang, L., Alexander, J., et al. (2020). Chikungunya Virus Strains from Each Genetic Clade Bind Sulfated Glycosaminoglycans as Attachment Factors. J. Virol. 94, e01500–20. https://doi.org/10.1128/JVI.01500-20.

McIlroy, D., Hönemann, M., Nguyen, N.-K., Barbier, P., Peltier, C., Rodallec, A., Halary, F., Przyrowski, E., Liebert, U., Hourmant, M., et al. (2020). Persistent BK Polyomavirus Viruria Is Associated with Accumulation of VP1 Mutations and Neutralization Escape. Viruses 12, 824. https://doi.org/10.3390/v12080824.

Neu, U., Allen, S.A., Blaum, B.S., Liu, Y., Frank, M., Palma, A.S., Ströh, L.J., Feizi, T., Peters, T., Atwood, W.J., et al. (2013). A Structure-Guided Mutation in the Major Capsid Protein Retargets BK Polyomavirus. PLoS Pathog. 9, e1003688. https://doi.org/10.1371/journal.ppat.1003688.

Nickeleit, V., Singh, H.K., Randhawa, P., Drachenberg, C.B., Bhatnagar, R., Bracamonte, E., Chang, A., Chon, W.J., Dadhania, D., Davis, V.G., et al. (2018). The Banff Working Group Classification of Definitive Polyomavirus Nephropathy: Morphologic Definitions and Clinical Correlations. J. Am. Soc. Nephrol. 29, 680–693. https://doi.org/10.1681/ASN.2017050477.

Nickeleit, V., Singh, H.K., Dadhania, D., Cornea, V., El-Husseini, A., Castellanos, A., Davis, V.G., Waid, T., and Seshan, S.V. (2021). The 2018 Banff Working Group classification of definitive polyomavirus nephropathy: A multicenter validation study in the modern era. Am. J. Transplant. 21, 669–680. https://doi.org/10.1111/ajt.16189.

Nickeleit, V., Hirsch, H.H., Binet, I.F., Gudat, F., Prince, O., Dalquen, P., Thiel, G., and Mihatsch, M.J. Polyomavirus Infection of Renal Allograft Recipients: From Latent Infection to Manifest Disease. J Am Soc Nephrol 10..

Pastrana, D.V., Brennan, D.C., Çuburu, N., Storch, G.A., Viscidi, R.P., Randhawa, P.S., and Buck, C.B. (2012). Neutralization Serotyping of BK Polyomavirus Infection in Kidney Transplant Recipients. PLoS Pathog. 8, e1002650. https://doi.org/10.1371/journal.ppat.1002650.

Peretti, A., Geoghegan, E.M., Pastrana, D.V., Smola, S., Feld, P., Sauter, M., Lohse, S., Ramesh, M., Lim, E.S., Wang, D., et al. (2018). Characterization of BK Polyomaviruses from Kidney Transplant Recipients Suggests a Role for APOBEC3 in Driving In-Host Virus Evolution. Cell Host Microbe 23, 628–635.e7. https://doi.org/10.1016/j.chom.2018.04.005.

Ramos, E., Drachenberg, C.B., Papadimitriou, J.C., Hamze, O., Fink, J.C., Klassen, D.K., Drachenberg, R.C., Wiland, A., Wali, R., Cangro, C.B., et al. (2002). Clinical course of polyoma virus nephropathy in 67 renal transplant patients. J. Am. Soc. Nephrol. JASN 13, 2145–2151. https://doi.org/10.1097/01.asn.0000023435.07320.81.

Randhawa, P.S., Vats, A., Zygmunt, D., Swalsky, P., Scantlebury, V., Shapiro, R., and Finkelstein, S. (2002). Quantitation of viral DNA in renal allograft tissue from patients with BK virus nephropathy1: Transplantation 74, 485–488. https://doi.org/10.1097/00007890-200208270-00009.

Schowalter, R.M., Pastrana, D.V., and Buck, C.B. (2011). Glycosaminoglycans and Sialylated Glycans Sequentially Facilitate Merkel Cell Polyomavirus Infectious Entry. PLoS Pathog. 7, e1002161. https://doi.org/10.1371/journal.ppat.1002161.

Shinohara, T., Matsuda, M., Cheng, S.H., Marshall, J., Fujita, M., and Nagashima, K. (1993). BK virus infection of the human urinary tract. J. Med. Virol. 41, 301–305. https://doi.org/10.1002/jmv.1890410408.

Viscount, H.B., Eid, A.J., Espy, M.J., Griffin, M.D., Thomsen, K.M., Harmsen, W.S., Razonable, R.R., and Smith, T.F. (2007). Polyomavirus polymerase chain reaction as a surrogate marker of polyomavirus-associated nephropathy. Transplantation 84, 340–345. https://doi.org/10.1097/01.tp.0000275205.41078.51.

Winn, M.D., Ballard, C.C., Cowtan, K.D., Dodson, E.J., Emsley, P., Evans, P.R., Keegan, R.M., Krissinel, E.B., Leslie, A.G.W., McCoy, A., et al. (2011). Overview of the CCP4 suite and current developments. Acta Crystallogr. D Biol. Crystallogr. 67, 235–242. https://doi.org/10.1107/S0907444910045749.

