## Supplemental Table 3 for "Structural and functional analysis of natural capsid variants reveals sialic-acid independent entry of BK polyomavirus"

**Supplemental Table 3. Supplementary glycan microarray document based on MIRAGE Glycan Microarray guidelines (doi:10.3762/mirage.3).**

<table>
<tr><th>Classification</th><th>Guidelines</th></tr>
<tr><td colspan="2">1. Sample: Glycan Binding Sample</td></tr>
<tr><td>Description of Sample</td><td><u>Sample names</u>:<br>Wild type (WT) BKPyV VP1<br>Single mutants: E73A BKPyV VP1, E73Q BKPyV VP1<br>Triple mutant: A72V-E73Q-E82Q (VQQ) BKPyV VP1<br>Double mutant: K69N-E82Q (N-Q) BKPyV VP1<br><u>Origin</u>: recombinant<br><u>Method of preparation:</u><br>Please see the “Protein expression and purification” section under Materials and Methods in the main text.</td></tr>
<tr><td>Sample modifications</td><td>Not relevant.</td></tr>
<tr><td>Assay protocol</td><td>Microarray analyses were performed essentially as described (Liu et al., Methods Mol. Biol. 2012), for modifications of the protocol please see “Glycan microarray screening” under Materials and Methods section in the main text.</td></tr>
<tr><td colspan="2">2. Glycan Library</td></tr>
<tr><td>Glycan description for defined glycans</td><td>Two glycan microarrays were used, both containing sequence-defined lipid-linked oligosaccharide probes, glycolipids or neoglycolipids (NGLs).<br>1) The ‘Ganglioside-focused array’ contained 26 ganglioside related probes. These are a sub-set of an array containing 64 glycan probes (in-house designation ‘Neuro-Glycan Array Set 1’, which will be published elsewhere). The names and sequences of the 26 probes are given below.
<table>
<tr><th>Position[a]</th><th>Probe</th><th>Structure[b]</th></tr>
<tr><td>1</td><td>Asialo-GM2</td><td>GalNAcβ-4Galβ-4Glcβ-Cer</td></tr>
<tr><td>2</td><td>Asialo-GM1</td><td>Galβ-3GalNAcβ-4Galβ-4Glcβ-Cer</td></tr>
<tr><td>3</td><td>GM4</td><td>NeuAcα-3Galβ-Cer</td></tr>
<tr><td>4</td><td>Haematoside</td><td>NeuAcα-3Galβ-4Glcβ-Cer</td></tr>
<tr><td>5</td><td>GM3</td><td>NeuAcα-3Galβ-4Glcβ-Cer</td></tr>
<tr><td>6</td><td>GM3(Gc)</td><td>NeuGcα-3Galβ-4Glcβ-Cer</td></tr>
<tr><td>7</td><td>GM1</td><td>Galβ-3GalNAcβ-4Galβ-4Glcβ-Cer<br>|<br>NeuAcα-3</td></tr>
<tr><td>8</td><td>GM1(Gc)</td><td>Galβ-3GalNAcβ-4Galβ-4Glcβ-Cer<br>|<br>NeuGcα-3</td></tr>
<tr><td>9</td><td>GM1b</td><td>NeuAcα-3Galβ-3GalNAcβ-4Galβ-4Glcβ-Cer</td></tr>
<tr><td>10</td><td>GM2</td><td>GalNAcβ-4Galβ-4Glcβ-Cer<br>|<br>NeuAcα-3</td></tr>
<tr><td>11</td><td>GD3</td><td>NeuAcα-8NeuAcα-3Galβ-4Glcβ-Cer</td></tr>
<tr><td>12</td><td>GD2</td><td>GalNAcβ-4Galβ-4Glcβ-Cer<br>|<br>NeuAcα-8NeuAcα-3</td></tr>
</table></td></tr>
</table>

| | | |
|---|---|---|
| 13 | GD1a | NeuAcα-3Galβ-3GalNAcβ-4Galβ-4Glcβ-Cer<br>NeuAcα-3 |
| 14 | GD1b | Galβ-3GalNAcβ-4Galβ-4Glcβ-Cer<br>NeuAcα-8NeuAcα-3 |
| 15 | GalNAc-GD1a(Ac,Gc) | GalNAcβ-4Galβ-3GalNAcβ-4Galβ-4Glcβ-Cer<br>NeuGcα-3 NeuAcα-3<br>GalNAcβ-4Galβ-3GalNAcβ-4Galβ-4Glcβ-Cer<br>NeuAcα-3 NeuGcα-3 |
| 16 | GT1a | NeuAcα-8NeuAcα-3Galβ-3GalNAcβ-4Galβ-4Glcβ-Cer<br>NeuAcα-3 |
| 17 | GT1b | NeuAcα-3Galβ-3GalNAcβ-4Galβ-4Glcβ-Cer<br>NeuAcα-8NeuAcα-3 |
| 18 | GQ1b | NeuAcα-8NeuAcα-3Galβ-3GalNAcβ-4Galβ-4Glcβ-Cer<br>NeuAcα-8NeuAcα-3 |
| 19 | Asialo-GM1-Tetra | Galβ-3GalNAcβ-4Galβ-4Glc-DH |
| 20 | GM1-penta | Galβ-3GalNAcβ-4Galβ-4Glc-DH<br>NeuAcα-3 |
| 21 | GM1(Gc)-penta | Galβ-3GalNAcβ-4Galβ-4Glc-DH<br>NeuGcα-3 |
| 22 | GM1b-DH | NeuAcα-3Galβ-3GalNAcβ-4Galβ-4Glc-DH |
| 23 | GD1a-hexa | NeuAcα-3Galβ-3GalNAcβ-4Galβ-4Glc-DH<br>NeuAcα-3 |
| 24 | GD3-tetra | NeuAcα-8NeuAcα-3Galβ-4Glc-DH |
| 25 | GD1b-DH | Galβ-3GalNAcβ-4Galβ-4Glc-DH<br>NeuAcα-8NeuAcα-3 |
| 26 | GT1c-DH | Galβ-3GalNAcβ-4Galβ-4Glc-DH<br>NeuAcα-8NeuAcα-8NeuAcα-3 |

[a] Probe position in the histogram charts (**Figure 2A**)

[b] Cer, ceramide; DH, 1,2-dihexadecyl-*sn*-glycero-3-phosphoethanolamine (DHPE).

2) A broad spectrum screening microarray contained 672 sequence-defined oligosaccharide probes **(Supplemental Table S2)**. These are a sub-set of a recently generated large screening microarray containing around 900 glycan probes (in-house designation 'Array Sets 42-56', which will be published elsewhere). The NGL probes are from the collection assembled in the course of research in the Glycosciences Laboratory

(https://glycosciences.med.ic.ac.uk/glycanLibraryList.html).

| | |
|---|---|
| Glycan description for undefined glycans | Not relevant. |
| Glycan modifications | No modification was carried out for natural glycolipids.<br>For NGLs, unless otherwise specified these were prepared from reducing oligosaccharides by reductive amination with the amino lipid, 1,2-dihexadecyl-*sn*-glycero-3-phosphoethanolamine [(DHPE) (Chai et al., Methods Enzymol. 2003)]; AO, NGLs prepared from reducing oligosaccharides by oxime ligation with an aminooxy functionalized DHPE [(AOPE) (Liu et al., Chem. Biol. 2007)]. |

| | |
|---|---|
| | For full description on the definition of lipid moieties of the glycan probes please see https://glycosciences.med.ic.ac.uk/docs/lipids.pdf. |
| **3. Printing Surface; e.g., Microarray Slide** | |
| Description of surface | Nitrocellulose-coated glass microarray slides. |
| Manufacturer | 16-pad UniSart® 3D Microarray Slide from Sartorius (Goettingen, Germany) |
| Custom preparation of surface | Not relevant. |
| Non-covalent Immobilisation | The lipid-linked oligosaccharide probes were formulated as liposomes by adding carrier lipids, 1,2-dihexanoyl-*sn*-glycero-3-phosphocholine (DHPC) and cholesterol for arraying and non-covalent immobilization on nitrocellulose-coated glass slides (Liu et al., Methods Mol. Biol. 2012). |
| **4. Arrayer (Printer)** | |
| Description of Arrayer | Nano-Plotter 2.1 (GeSiM, Radeberg, Germany). |
| Dispensing mechanism | Non-contact liquid delivery with four dispensing tips. |
| Glycan deposition | Approximately 0.33 nl was printed per spot.<br>Lipid-linked glycan probes were printed at 2 and 5 fmol per spot in duplicate. |
| Printing conditions | The printing solutions were all aqueous based. Printing was performed at ambient temperature and relative humidity of 58%.<br>The 'liposome' printing solutions contained 100 pmol/µl of DHPC and cholesterol (both from SIGMA) as lipid carriers in addition to the lipid-linked glycan probes. The concentrations of the lipid-linked glycan probes were 5 and 15 pmol/µl for the 2 and 5 fmol per spot levels, respectively.<br>The printing solutions also contained Cy3 NHS ester (GE Healthcare) at 20 ng/ml (26 fmol/µl) as a marker to monitor the printing process. |
| **5. Glycan Microarray with "Map"** | |
| Array layout | The 'Ganglioside-focused array' was printed in 16-pad format. Each array slide contained 16-pad subarrays. Each pad was set up for printing 64 probes maximum, each at 2 levels in duplicate (four spots for one probe in a row); up to 256 spots (16x16) in total in each pad.<br>The 672 lipid-linked probes in the screening arrays were printed on multiple subarrays for parallel binding analyses. |
| Glycan identification | The 'Ganglioside-focused array' was analysed with cholera toxin and a number of polyomavirus VP1 proteins with known specificities, e.g. simian virus 40 VP1 (Campanero-Rhodes et al., J Virol. 2007) for quality control purposes. |

| | |
|---|---|
| and quality control | The quality control of the screening microarrays of sequence-defined glycan probes was carried out with: biotinylated plant lectins - *Ricinus Communis* Agglutinin I ($RCA_{120}$), *Aleuria aurantia* lectin (AAL), Concanavalin A (ConA) and WGA (Vector Laboratories), a wide range of anti-carbohydrate antibodies, and a number of viral adhesive proteins that we have published previously.<br>These data will be described elsewhere and are available upon request. |
| **6. Detector and Data Processing** | |
| Scanning hardware | GenePix 4300A (Molecular Devices, UK) |
| Scanner settings | Scanning resolution: 10 µm / pixel<br>Laser channel: Red (scan wavelength 635 nm)<br>PMT: 350<br>Scan power: 100% to achieve maximum signal without spot saturation. |
| Image analysis software | GenePix® Pro 7 (Molecular Devices) |
| Data processing | The gpr files were entered into an in-house microarray database using software (designed by Mark Stoll, http://www.beilstein-institut.de/en/publications/proceedings/glyco-2009) for data processing. No particular normalization method or statistical analysis was used for the results of the screening arrays. |
| **7. Glycan Microarray Data Presentation** | |
| Data presentation | The microarray binding results are in **Figure 2, Supplemental Figure 2** and **Supplemental Table 2.** |
| **8. Interpretation and Conclusion from Microarray Data** | |
| Data interpretation | No software or algorithms were used to interpret processed data. |
| Conclusions | The wild type BKPyV VP1 and its E73A, E73Q and VQQ mutants bound to sialylated glycan probes in the arrays with different binding profiles and fluorescence intensities. No significant binding was detected with the N-Q mutant VP1. |
